# Resting-State Prefrontal Connectivity Varies With Placebo-Induced Hunger Expectations in Healthy Women

**DOI:** 10.64898/2026.09.03.749135

**Authors:** Iraj Khalid, Benjamin Flament, Belina Rodrigues, Jason Morris, Phillipe Fossati, Karin Meissner, Liane Schmidt

## Abstract

Placebo effects, in which a patient’s symptoms and subjective well-being improve in response to an inactive treatment, medication, or intervention (a placebo), are well known in clinical research. Basic research in human cognition and neuroscience has shown that these effects are linked to prognostic expectations of effectiveness and to recruitment of prefrontal cortex regions, including the medial (mPFC), ventromedial (vmPFC), and dorsolateral prefrontal cortex (dlPFC). However, how these expectations influence intrinsic resting-state brain connectivity remains unclear. Here, we conducted seed-based resting-state functional connectivity analyses in 60 healthy, fasting female participants undergoing a placebo intervention that induced hunger expectations. In line with previously published results, the intervention induced hunger expectations and biased hunger sensations. We found that decreased hunger expectations attenuated mPFC-angular gyrus connectivity within the default mode network, while increased hunger expectations reduced vmPFC-dlPFC connectivity and enhanced dlPFC-thalamus coupling within frontoparietal and interoceptive networks. These results suggest that placebo-induced hunger expectations modulate prefrontal cortex connections with regions involved in valuation, cognitive control, and interoception. This modulation may reflect neural mechanisms by which beliefs shape bodily sensations under placebo conditions, advancing understanding of the neural substrates of expectation-driven sensory experiences.

**Highlights:**

◻ Decreased-hunger expectations attenuate mPFC–angular gyrus connectivity.
◻ Increased-hunger expectations attenuate vmPFC–dlPFC connectivity.
◻ Increased-hunger expectations strengthen dlPFC–thalamus connectivity.

## INTRODUCTION

Placebo effects are well known from clinical trials, where they serve as controls for effects not attributable to the specific treatment itself (i.e., non-specific effects). These effects occur when patients experience improvements in symptoms and subjective well-being after receiving a sham treatment—such as a pharmacologically inactive substance or a procedure that mimics the active treatment but lacks its active components (Schmidt and Koban, 2022). Importantly, research in cognitive neuroscience has shown that placebo effects are mediated by active physiological and psychological processes. Notably, prognostic beliefs about a treatment’s effectiveness, known as outcome and response expectations, are powerful psychological factors that can bias perception and even override sensory evidence (Colloca and Miller, 2011; Finan and Colloca, 2025). A large body of brain imaging studies in the pain domain has shown that expectation-based placebo effects are associated with reduced activation of nociceptive brain regions, including the dorsal anterior cingulate cortex (dACC), anterior insula, and thalamus (for review see: Ashar et al., 2017; Wager and Atlas, 2015; Zunhammer et al., 2021). At the same time, placebo cues consistently activate non-nociceptive brain regions such as the medial (mPFC), ventromedial (vmPFC), and dorsolateral prefrontal cortex (dlPFC) (Wager and Atlas, 2015). These regions are also engaged during placebo effects in other, less aversive domains of behavior and judgment, including learning, taste pleasantness, hunger sensations, and food choices (I. Khalid et al., 2024; Plassmann et al., 2008; Schmidt et al., 2017, 2014).

In more detail, the mPFC has been found to encode expectations about a placebo’s effect on hunger sensations at the time of food choice (Khalid et al., 2024), which aligns with its role in encoding expected pain during placebo analgesia (Ashar et al., 2017; Jepma et al., 2018; Wager and Atlas, 2015; Zunhammer et al., 2021). The vmPFC, by contrast, encoded the subjective value of food based on hunger expectations (Khalid et al., 2024), aligning with its role in assigning affective meaning to specific contexts, such as those created by placebo interventions (Roy et al., 2012). The dlPFC is involved in resolving interference and has been shown to mediate the effects of hunger suggestions on hunger sensations (Khalid et al., 2024). This is consistent with its well-known role in cognitive self-regulation during decision-making (Rudorf and Hare, 2014). These findings come from task-based fMRI, which measures the relationship between explicit behavioral responses (such as ratings and choices) and the brain’s blood oxygenation level-dependent (BOLD) signal. However, it remains unclear whether expectations induced by a placebo intervention also alter more stable, intrinsic characteristics of the BOLD signal. Resting-state fMRI (RS-fMRI) offers a non-invasive method to address this question by assessing spontaneous fluctuations in the BOLD signal when no explicit task is performed (Biswal and Uddin, 2025). RS-fMRI can reveal patterns of functional connectivity—spatiotemporal correlations between brain regions—that reflect the brain’s intrinsic functional organization, which can vary with psychological states and clinical conditions (Buckner et al., 2008). Understanding how such contexts influence these spontaneous neural fluctuations may provide insight into the neural mechanisms underlying under-defined problems such as placebo effects.

To our knowledge, only a handful of studies have examined resting-state brain activity following placebo interventions targeting pain (Kong et al., 2013; Spanou et al., 2026; Tétreault et al., 2016a), depression (Gerlach et al., 2026), or Parkinson’s disease (Joineau et al., 2026). Findings from the pain domain indicate that frontoparietal cortex regions exhibit stronger connectivity with the rostral anterior cingulate cortex (rACC) during placebo analgesia (Kong et al., 2013). Additionally, individual differences in resting-state functional connectivity of the right midfrontal gyrus, a region within the dlPFC, have been shown to distinguish placebo responders from non-responders and to predict the magnitude of placebo analgesia (Trétreault et al. 2016a).

Here we aimed to provide evidence of how placebo-induced expectations about more appetitive domains of experiences and sensations influence resting-state brain connectivity in healthy participants. We combined a placebo intervention targeting hunger sensations with RS-fMRI in healthy female participants in a fasted state. We focused on three seed regions of interest—the mPFC, vmPFC, and dlPFC—which, in the same sample, have previously been shown to underpin an ’appetitive’ placebo effect on hunger sensations and dietary decision-making (Khalid et al., 2024). We examined the functional connectivity of these three regions with the rest of the brain and tested the hypothesis that this connectivity would be moderated by placebo-induced hunger expectations. Specifically, we used seed-to-voxel whole-brain connectivity analyses to assess how (1) each seed ROI connected to the rest of the brain irrespective of placebo-induced hunger expectations, and how these connectivity patterns (2) varied with the strength of placebo-induced prognostic hunger expectations and (3) differed between participants who expected a decrease versus an increase in hunger.

## RESULTS

### Placebo effects on hunger expectations and sensations

The placebo intervention induced prognostic beliefs about the drink’s effect on hunger, including how much it was expected to increase or decrease hunger on a scale from 1 to 10 (mean_decreased_ ± SEM = 5.5 ± 0.3; mean_increased_ ± SEM = 5.3 ± 0.35). There was no significant difference between the hunger-suggestion groups (t(58) = 0.45, p = .65, Cohen’s d = 0.116, 95% CI [–0.39 to 0.62]), as shown by a two-tailed, two-sample t-test (Fig. 1a). When referencing the expectation ratings in each group to one, which on the rating scale indicated minimum expected effectiveness, both groups were significantly expecting more than minimum effectiveness (mean_centered_decreased_ ± sem = 4.5 ± 0.3, t(29) = 16.2, p < 0.001, Cohen’s d = 2.9, 95 % CI [2.1 to 3.8]; mean_centered_increased_ ± sem = 4.3 ± 0.35; t(29) = 12.4, p <0.001, Cohen’s d = 0.34, 95%CI [1.6 to 2.9]).

**Figure 1.**
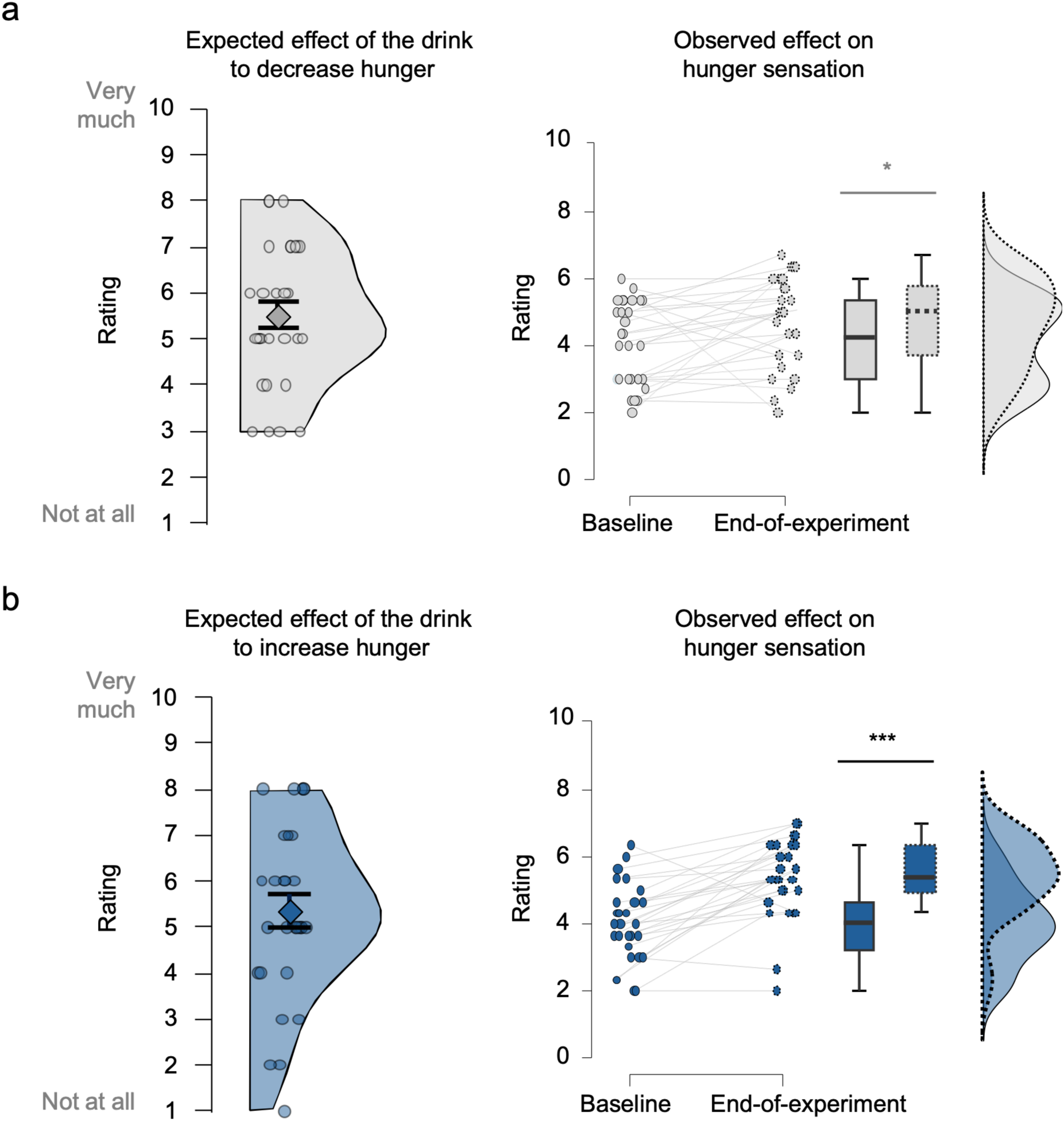
Expectancy across groups in the full RS-fMRI sample (n=60) and hunger ratings across groups in hungry-at-baseline subsample (n=56). **a.** For the decreased hunger suggestion group. b. For the increased hunger suggestion group. The expectancy ratings were collected in response to the question: How much do you expect the drink to decrease/increase your hunger? The hunger ratings were averaged across three ratings: hedonic (How pleasant would it be to eat right now?), homeostatic (How much could you eat right now?), and subjective general hunger (How hungry are you right now?). Boxplot graphs display the 95% confidence intervals, with boxes indicating the interquartile range from Q1 (25th percentile) to Q3 (75th percentile). The black horizontal lines indicate medians, and the whiskers range from minimum to maximum values and span 1.5 times the interquartile range. The dots beside each box plot correspond to individual participants. *p < 0.05, *** p<0.001 paired, two-tailed t-test.

In line with previously reported results (Khalid et al. 2024) the hunger sensations reported at baseline, in the hungry-at-baseline only participants (n=56) increased from the start to the end of the experiment (main effect of time (baseline-to-end): ß = 0.49, SE = 0.09, p< 0.001), but the increase was steeper in the increased compared to the decreased hunger suggestion group (group by time: ß = 0.2, SE = 0.09, p = 0.033, Fig. 1b). Note, the interaction between group and time was also significant when comparing baseline hunger ratings to hunger ratings given immediately after the administration of the placebo (group by baseline-to-post: ß = 0.17, SE = 0.08, p = 0.35), but not when comparing post-placebo administration to end of the experiment ratings (group by post-to-end: ß = 0.02, SE = 0.09, p = 0.87).

These results align with previously published findings (Hoffmann et al., 2018; Khalid et al., 2024) and support the effectiveness of the placebo intervention in inducing expectations and biasing hunger sensations in the albeit smaller RS-fMRI participant sample.

### Seed-to-voxel RS-fMRI connectivity results in RS-fMRI full sample (n=60)

Our main hypothesis focused on the seed-to-voxel resting-state connectivity of three regions of interest (ROIs): the mPFC, vmPFC, and dlPFC. We examined (1) the overall functional connectivity of each seed with the rest of the brain, (2) how this connectivity was moderated by prognostic hunger expectations within each suggestion group, and (3) how specifically the moderation by hunger expectations differed between the decreased and increased hunger suggestion groups.

#### (1) Overall seed-based resting-state connectivity

At a conservative false discovery rate (FDR) corrected voxel threshold of p_FDR_<0.05 and a family-wise error-corrected peak threshold of p_FWE_<0.05 the mPFC seed (MNI = [2, 58, 22]) connected positively with clusters in the default mode and limbic networks (Fig 2). These clusters were located mainly around the mPFC seed with peaks in the ventromedial and posterior cingulate cortex, as well as in the middle temporal gyrus. The mPFC seed connected negatively with a dorsolateral prefrontal cortex cluster located within the frontoparietal resting-state network (Fig. 2). At a more lenient peak threshold of p_FWE_<0.05 on the cluster level, the dlPFC decoupling became bilateral, and two more prominent bilateral clusters emerged located on the angular gyrus (Fig. 2, Table 1).

**Figure 2.**
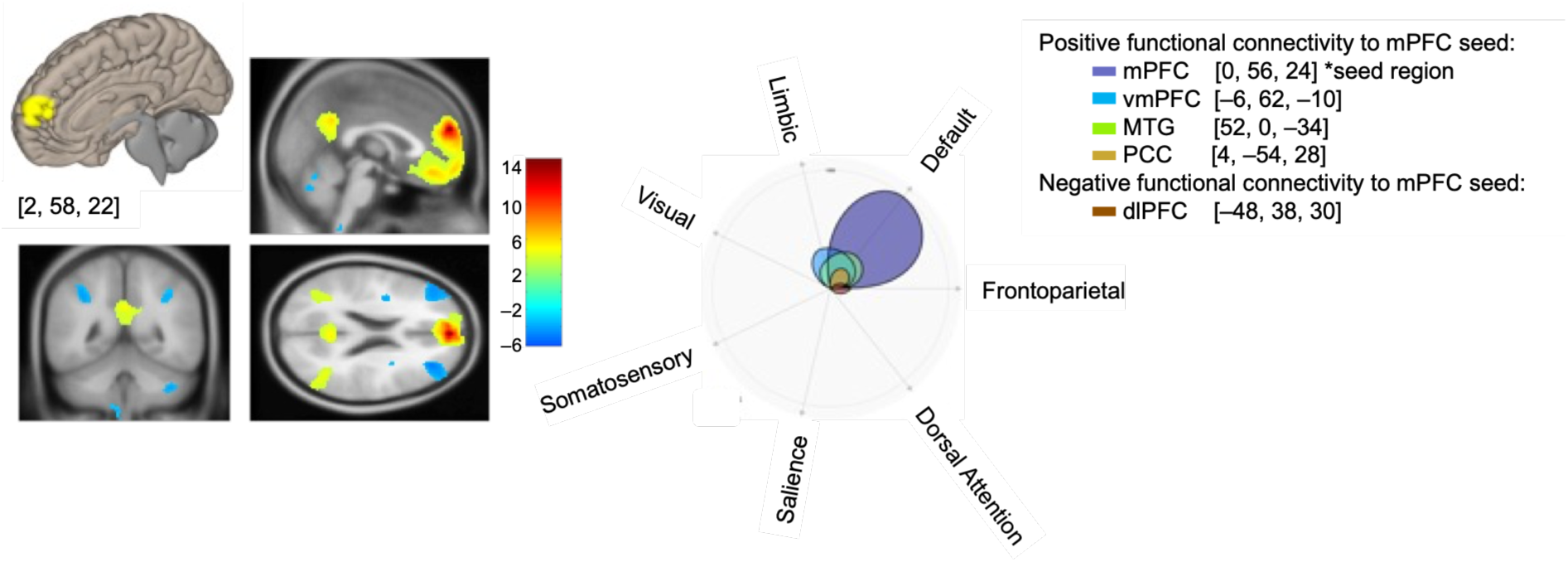
mPFC–to-voxel connectivity overall (RS-FMRI full sample, n=60). Statistical parametric maps (SPMs) show voxels that covaried positively (in red and yellow) and negatively (in blue) with the mPFC seed region, which is shown in yellow as 5-mm-radius spheres centered around the corresponding Montreal Neurological Institute (MNI) [x, y, z] coordinates and superimposed on the 3-D anatomical brains. SPMs are displayed for visualization at a whole-brain height threshold of p < 0.001 uncorrected and superimposed on the average anatomical brain. Polar plots localize clusters and cluster sizes (k voxels) within the CONN toolbox’s custom-built resting-state network atlas. They are thresholded at a voxel threshold of pFDR < 0.05 (false discovery rate corrected) and a peak threshold of pFWE < 0.05 (family-wise error corrected).

**Table 1:**
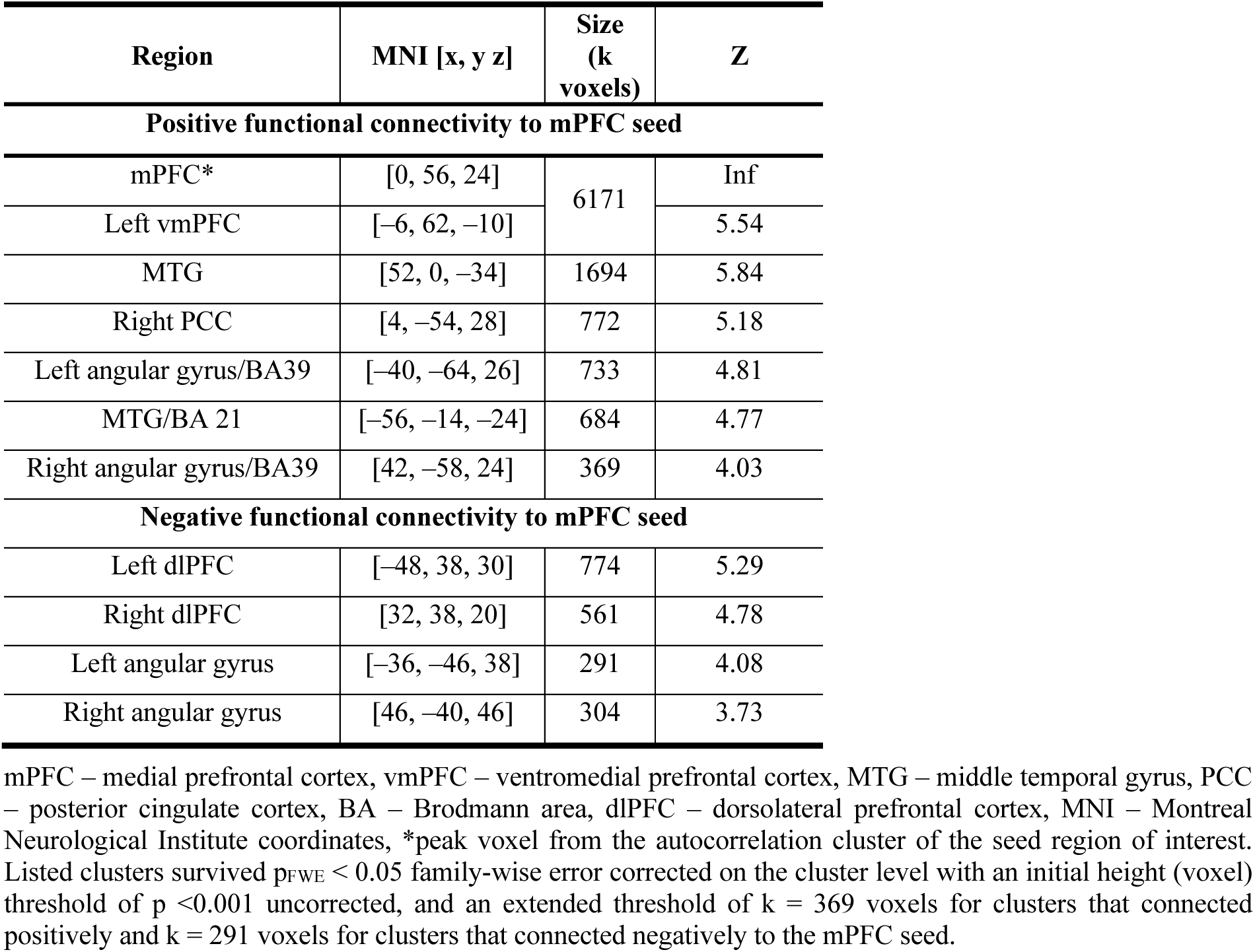
Number of voxels located in canonical resting-state networks for the mPFC seed-to-voxel connectivity (RS-fMRI full sample, n=60)

| Region | MNI [x, y z] | Size (k voxels) | Z |
| --- | --- | --- | --- |
| **Positive functional connectivity to mPFC seed** |  |  |  |
| mPFC* | [0, 56, 24] | 6171 | Inf |
| Left vmPFC | [-6, 62, -10] |  | 5.54 |
| MTG | [52, 0, -34] | 1694 | 5.84 |
| Right PCC | [4, -54, 28] | 772 | 5.18 |
| Left angular gyrus/BA39 | [-40, -64, 26] | 733 | 4.81 |
| MTG/BA 21 | [-56, -14, -24] | 684 | 4.77 |
| Right angular gyrus/BA39 | [42, -58, 24] | 369 | 4.03 |
| **Negative functional connectivity to mPFC seed** |  |  |  |
| Left dlPFC | [-48, 38, 30] | 774 | 5.29 |
| Right dlPFC | [32, 38, 20] | 561 | 4.78 |
| Left angular gyrus | [-36, -46, 38] | 291 | 4.08 |
| Right angular gyrus | [46, -40, 46] | 304 | 3.73 |

mPFC – medial prefrontal cortex, vmPFC – ventromedial prefrontal cortex, MTG – middle temporal gyrus, PCC – posterior cingulate cortex, BA – Brodmann area, dlPFC – dorsolateral prefrontal cortex, MNI – Montreal Neurological Institute coordinates, \*peak voxel from the autocorrelation cluster of the seed region of interest. Listed clusters survived $p_{FWE} < 0.05$ family-wise error corrected on the cluster level with an initial height (voxel) threshold of $p < 0.001$ uncorrected, and an extended threshold of $k = 369$ voxels for clusters that connected positively and $k = 291$ voxels for clusters that connected negatively to the mPFC seed.

At a conservative false discovery rate (FDR) corrected voxel threshold of p_FDR_<0.05 and a family-wise error-corrected peak threshold of p_FWE_<0.05 the vmPFC seed (MNI = [0, 52, –12]) covaried positively with clusters in the default mode, limbic, visual, frontoparietal, and dorsal attention networks (Fig. 3, Table 2). These clusters were in the left posterior cingulate cortex, the left dorsomedial prefrontal cortex, and temporal and parietal regions, such as the left middle temporal gyrus, which extended further into the angular gyrus/precuneus and the bilateral parahippocampus (Fig. 3). Note that activations became mostly bilateral at a more lenient, albeit corrected, threshold of p_FWE_ <0.05 on the cluster level with an initial height (voxel) threshold of p < 0.001 uncorrected, extend threshold k =260 voxels, the vmPFC connected negatively to regions located in the dorsolateral, ventrolateral, and inferior parietal cortex (Table 2).

**Figure 3.**
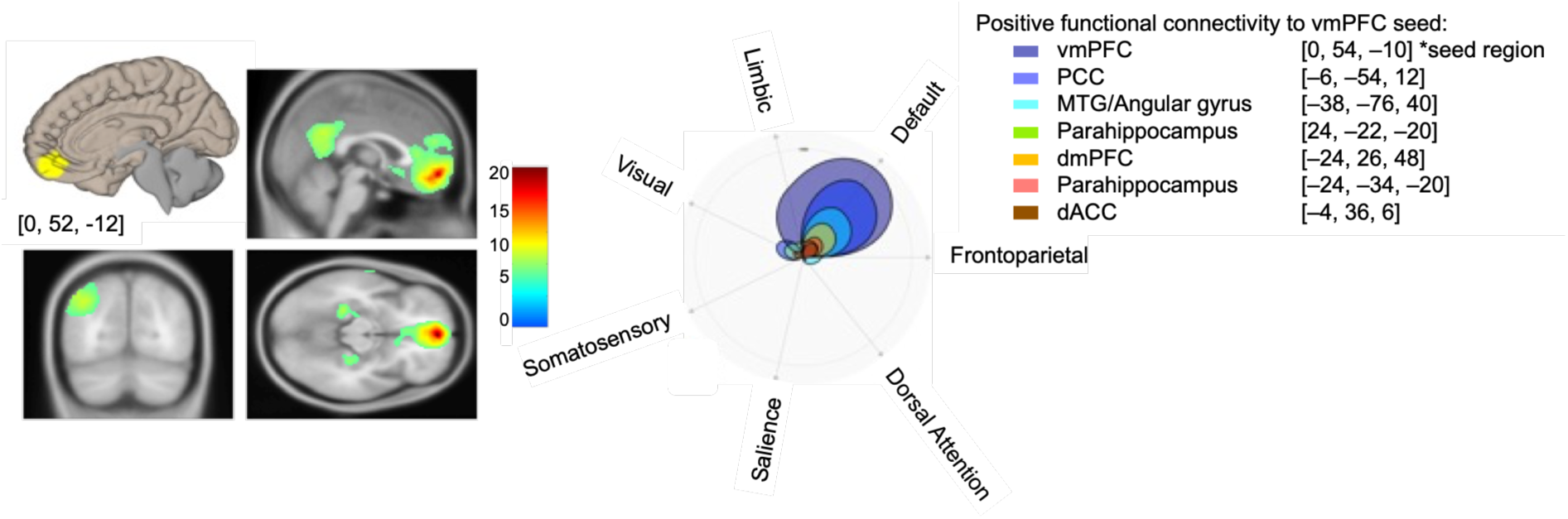
vmPFC–to-voxel connectivity overall (RS-FMRI full sample, n=60). Statistical parametric maps (SPMs) show voxels that covaried positively (red and yellow) with the vmPFC seed region, shown in yellow as 5-mm-radius spheres centered on the corresponding Montreal Neurological Institute (MNI) [x, y, z] coordinates and superimposed on the 3-D anatomical brains. SPMs are displayed for visualization at a whole-brain height threshold of p < 0.001 uncorrected and superimposed on the average anatomical brain. Polar plots localize clusters and cluster sizes (k voxels) within the CONN toolbox’s custom-built resting-state network atlas. They are thresholded at a voxel threshold of pFDR < 0.05 (false discovery rate corrected) and a peak threshold of pFWE < 0.05 (family-wise error corrected).

**Table 2:**
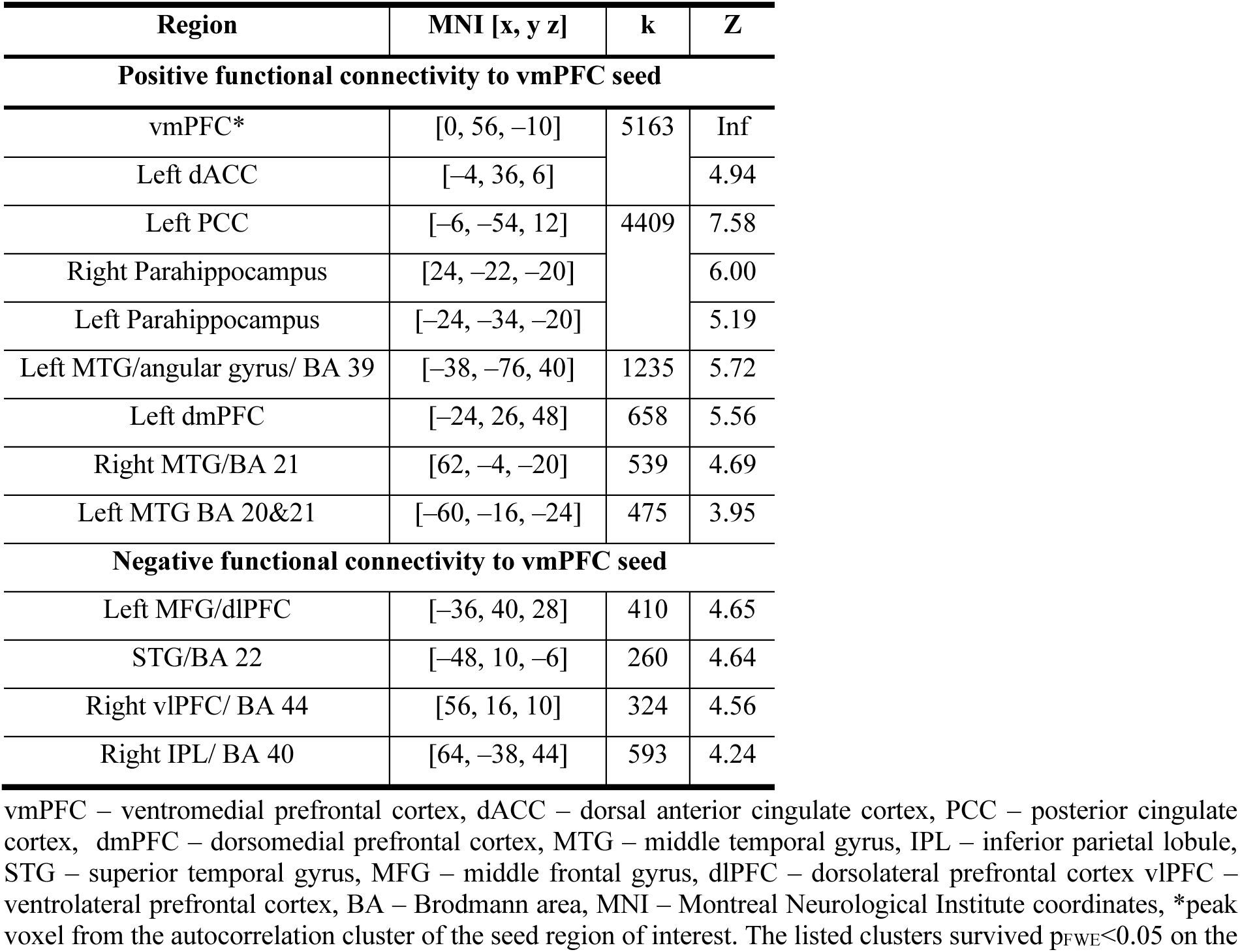
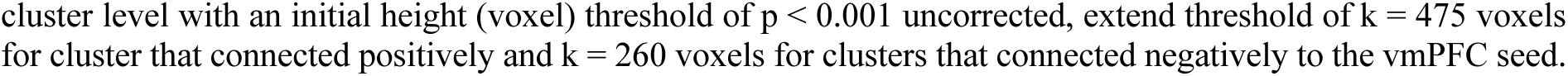
Number of voxels located in canonical resting state networks for the vmPFC seed-to-voxel connectivity (RS-fMRI full sample, n=60).

| Region | MNI [x, y, z] | k | Z |
| --- | --- | --- | --- |
| **Positive functional connectivity to vmPFC seed** |  |  |  |
| vmPFC* | [0, 56, -10] | 5163 | Inf |
| Left dACC | [-4, 36, 6] |  | 4.94 |
| Left PCC | [-6, -54, 12] | 4409 | 7.58 |
| Right Parahippocampus | [24, -22, -20] |  | 6.00 |
| Left Parahippocampus | [-24, -34, -20] |  | 5.19 |
| Left MTG/angular gyrus/ BA 39 | [-38, -76, 40] | 1235 | 5.72 |
| Left dmPFC | [-24, 26, 48] | 658 | 5.56 |
| Right MTG/BA 21 | [62, -4, -20] | 539 | 4.69 |
| Left MTG BA 20&21 | [-60, -16, -24] | 475 | 3.95 |
| **Negative functional connectivity to vmPFC seed** |  |  |  |
| Left MFG/dlPFC | [-36, 40, 28] | 410 | 4.65 |
| STG/BA 22 | [-48, 10, -6] | 260 | 4.64 |
| Right vlPFC/ BA 44 | [56, 16, 10] | 324 | 4.56 |
| Right IPL/ BA 40 | [64, -38, 44] | 593 | 4.24 |

vmPFC – ventromedial prefrontal cortex, dACC – dorsal anterior cingulate cortex, PCC – posterior cingulate cortex, dmPFC – dorsomedial prefrontal cortex, MTG – middle temporal gyrus, IPL – inferior parietal lobule, STG – superior temporal gyrus, MFG – middle frontal gyrus, dlPFC – dorsolateral prefrontal cortex vlPFC – ventrolateral prefrontal cortex, BA – Brodmann area, MNI – Montreal Neurological Institute coordinates, \*peak voxel from the autocorrelation cluster of the seed region of interest. The listed clusters survived $p_{FWE} < 0.05$ on the
cluster level with an initial height (voxel) threshold of $p < 0.001$ uncorrected, extend threshold of $k = 475$ voxels for cluster that connected positively and $k = 260$ voxels for clusters that connected negatively to the vmPFC seed.

The dlPFC seed (MNI = [40, 42, 26]) covaried with clusters of the frontoparietal and salience networks (Fig. 4) at a conservative false discovery rate (FDR) corrected voxel threshold of p_FDR_<0.05 and a family-wise error-corrected peak threshold of p_FWE_<0.05. These clusters were in the contralateral left dlPFC and right middle frontal gyrus (Fig. 4, Table 3). Negative functional connectivity was observed at a more lenient cluster-corrected threshold of p_FWE_ < 0.05, initial height (voxel) threshold of p<0.001 uncorrected, extend threshold of k = 370 voxels in the middle temporal gyrus and left superior frontal gyrus, Brodmann area 9 (Table 3).

**Figure 4.**
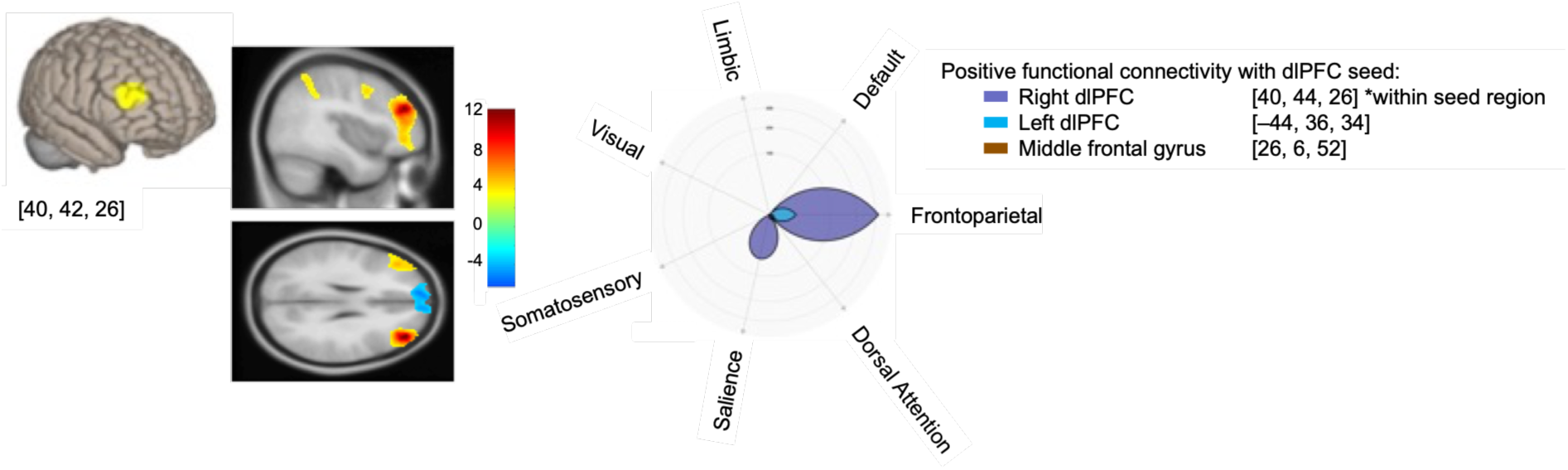
dlPFC–to-voxel connectivity overall (RS-FMRI full sample, n=60). Statistical parametric maps (SPMs) show voxels that covaried positively (red and yellow) and negatively (blue) with the right dlPFC seed region, shown in yellow as 5-mm-radius spheres centered on the corresponding Montreal Neurological Institute (MNI) [x, y, z] coordinates and superimposed on the 3-D anatomical brains. SPMs are displayed for visualization at a whole-brain height threshold of p < 0.001 uncorrected and superimposed on the average anatomical brain. Polar plots localize clusters and cluster sizes (k voxels) within the CONN toolbox’s custom-built resting-state network atlas. They are thresholded at a voxel threshold of pFDR < 0.05 (false discovery rate corrected) and a peak threshold of pFWE < 0.05 (family-wise error corrected).

**Table 3:** Number of voxels located in canonical resting state networks for the dlPFC seed-to-voxel connectivity (RS-fMRI full sample, n=60)

| Region | MNI [x, y z] | k | Z |
| --- | --- | --- | --- |
| Positive functional connectivity with dlPFC seed |  |  |  |
| Right dlPFC* | [40, 44, 26] | 1603 | Inf |
| Left dlPFC | [-44, 36, 34] | 674 | 5.76 |
| Right MFG | [26, 6, 52] | 438 | 4.99 |
| IFL/BA 40 | [52, -40, 44] | 704 | 4.48 |
| Negative functional connectivity with dlPFC seed |  |  |  |
| MTG | [-46, -28, -18] | 370 | 5.38 |
| Left MFG/SFG/BA 9 | [-10, 58, 28] | 1474 | 4.73 |

dlPFC – dorsolateral prefrontal cortex, MFG – Middle frontal gyrus, IFL – inferior parietal lobule, MTG – middle temporal gyrus, SFG – superior frontal gyrus, BA – Brodmann area, MNI – Montreal Neurological Institute coordinates, \*peak voxel from the autocorrelation cluster of the seed region of interest. The listed clusters survived $p_{FWE} < 0.05$ on the cluster level with an initial height (voxel) threshold of $p < 0.001$ uncorrected, extend threshold of $k = 438$ voxels for clusters that connected positively, and $k = 370$ voxels for clusters that connected negatively to the dlPFC seed.

#### (2a) Moderation of seed-to-voxel resting state connectivity by decreased hunger expectations

Expectations about the drink’s effectiveness in decreasing hunger negatively moderated the functional connectivity between the mPFC seed and the left parietal cortex region encompassing the precuneus (Brodmann area 39) and angular gyrus (MNI = [-34, -70, 28], p_FWE_ <0.05, whole-brain family-wise error corrected at the peak and cluster level, Fig. 5a, b). The negative moderation of the mPFC – angular gyrus coupling by decreased hunger expectations was significantly stronger compared to the increased hunger suggestion group (r_decreased_ = –0.59, p < 0.001 versus r_increased_ = –0.10, p = 0.58, z = –2.12, p = 0.034, two-tailed, Fisher r-to-z transformation test, Fig. 5c).

**Figure 5.**
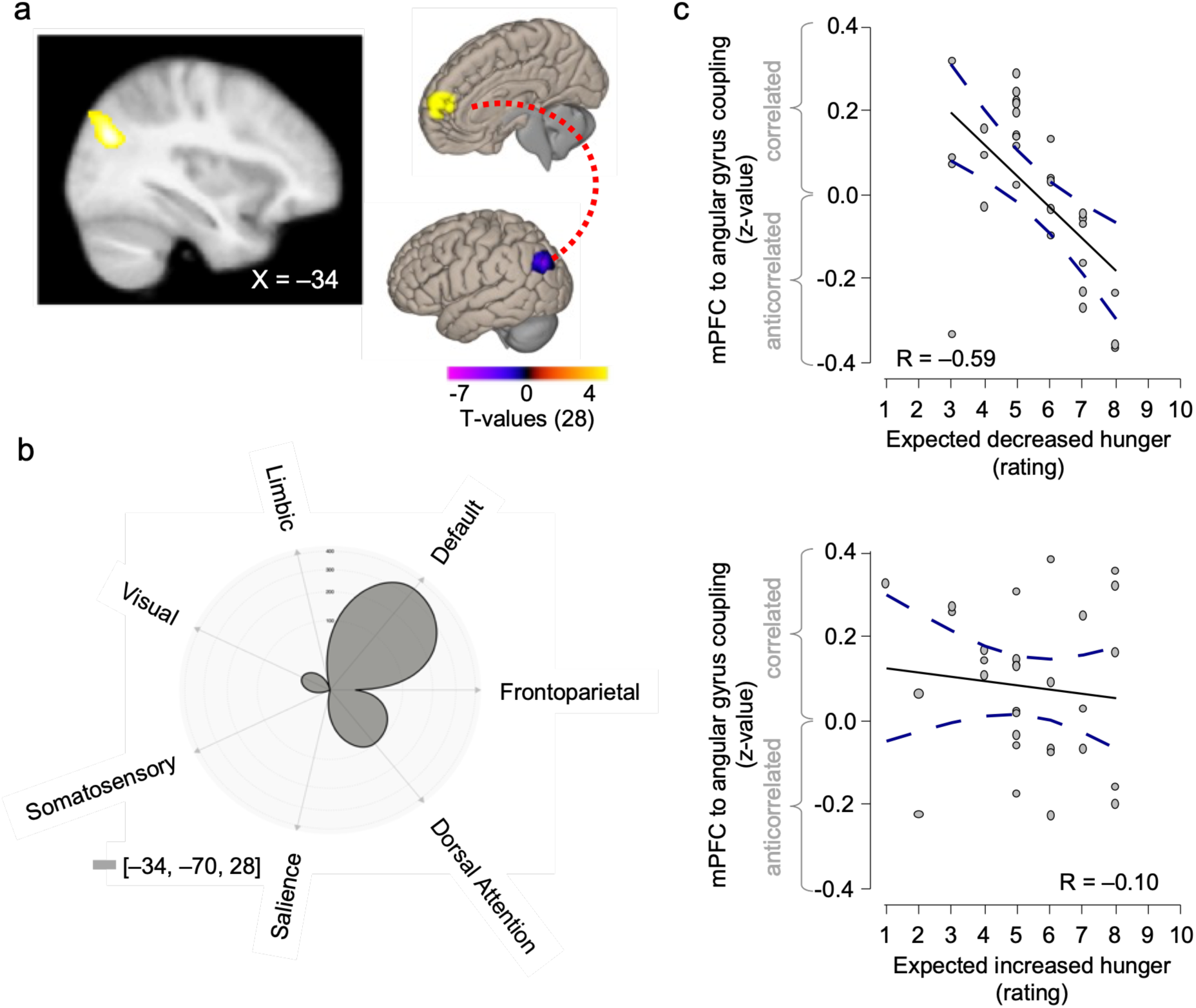
Effects of decreased hunger expectancies on mPFC-to-brain connectivity (RS-fMRI full subsample of n=30/60). (a) The statistical parametric maps (SPMs) and schematic 3D illustration show significant voxels in orange and yellow that survived pFWE<0.05 whole-brain family-wise error correction at the peak and cluster level, and are displayed at a whole-brain, height (voxel) threshold of p<0.001 uncorrected. (b) The polar plot shows the distribution of significant voxels across resting-state networks at pFWE<0.05, whole-brain cluster-level family-wise error correction. (c) Scatterplots show individual participants’ hunger expectations and z-values extracted from angular gyrus individual peaks selected via a leave-one-out procedure. R – Pearson’s correlation coefficient.

This result indicated that participants with stronger expectations that the placebo would reduce their hunger exhibited greater decoupling of the mPFC from the angular gyrus. In contrast, decreased hunger expectations did not significantly affect the seed-to-voxel connectivity of the vmPFC or dlPFC.

#### (2b) Moderation of seed-to-voxel resting-state connectivity by increased hunger expectations

Increased hunger expectations negatively moderated vmPFC seed coupling to a region within the dorsomedial (dmPFC) boundaries of the dlPFC, encompassing Brodmann Area 46 (MNI=[34, 52, 10], p_FWE_ <0.05, whole-brain family-wise error corrected, Fig. 6a). These significant dmPFC voxels were exclusively located within the frontoparietal resting state network (k = 217 voxels; Fig. 6b). The effect of hunger expectations on vmPFC – dmPFC functional decoupling was non-significant in the decreased hunger suggestion group, and complementary evidence from the comparison of the absolute Pearson’s coefficients showed its moderation by hunger expectations was significantly more pronounced in the increased hunger suggestion group (r_decreased_= 0.053, p = 0.782 versus r_increased_ = –0.456, p = 0.011, z = – 1.99, p = 0.046; two-tailed, Fisher’s r-to-z transformation test, Fig. 6c).

**Figure 6.**
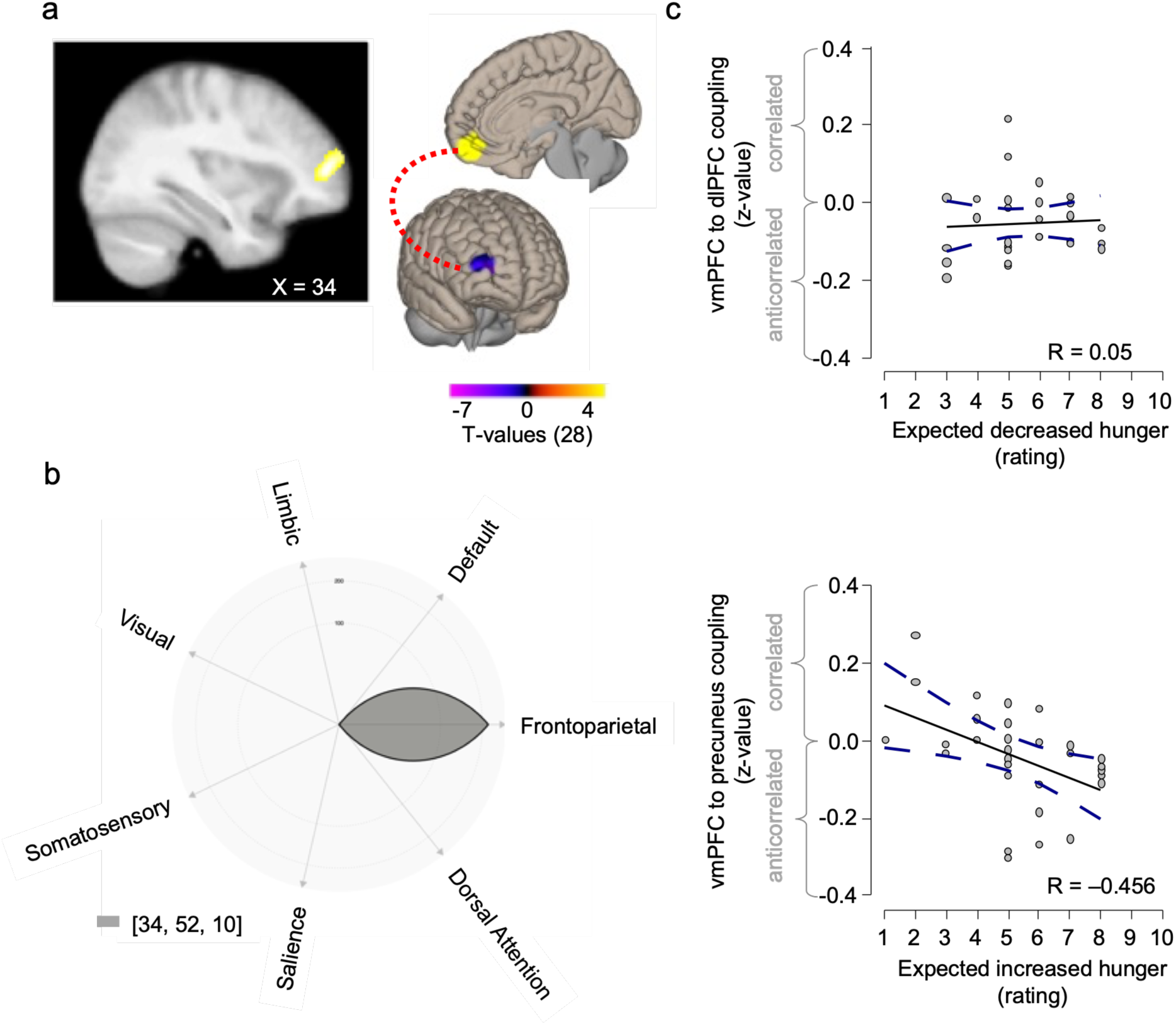
Effects of increased hunger expectancies on vmPFC-to-brain connectivity (RS-fMRI full subsample of n=30/60) (a) Statistical parametric maps (SPMs) and a schematic 3D-illustration show significant voxels in orange and yellow that survived pFWE < 0.05 whole-brain family-wise error correction at the cluster level and are displayed for visualization at a whole-brain, height (voxel) threshold of p < 0.001, uncorrected. (b) Polar plot showing the distribution of significant voxels across resting-state networks located solely within the frontoparietal network at pFWE < 0.05, whole-brain cluster-level family-wise error correction. (c) Scatterplots show individual participants’ hunger expectations and z-values extracted from individual peaks in the dlPFC selected via a leave-one-out procedure. R – Pearson’s correlation coefficient

Increased hunger expectations did not significantly moderate the seed-to-voxel connectivity of the two other seed ROIs – the mPFC and the vmPFC.

#### (3) Seed-to-voxel resting-state connectivity moderation by increased versus decreased hunger expectations

In direct comparison, the influence of hunger expectations on seed-to-voxel functional connectivity revealed that increased versus decreased hunger expectations were linked to a positive functional connection between the dlPFC seed region and the thalamus (MNI = [16, - 28, 4]; p_FWE_ < 0.05 whole-brain family-wise error corrected at the cluster level, Fig. 7a). This positive association was stronger for increased than for decreased hunger expectations, as suggested by complementary evidence from direct two-sampled t-tests of SPMs (Fig. 7a) and a significant difference in absolute Pearson’s coefficients (r_decreased_ = –0.279, p = 0.136 versus r_increased_ = 0.426, p = 0.019; z = -2.72, p = 0.0065; two-tailed, Fisher’s r-to-z transformation test, Fig. 7b).

**Figure 7.**
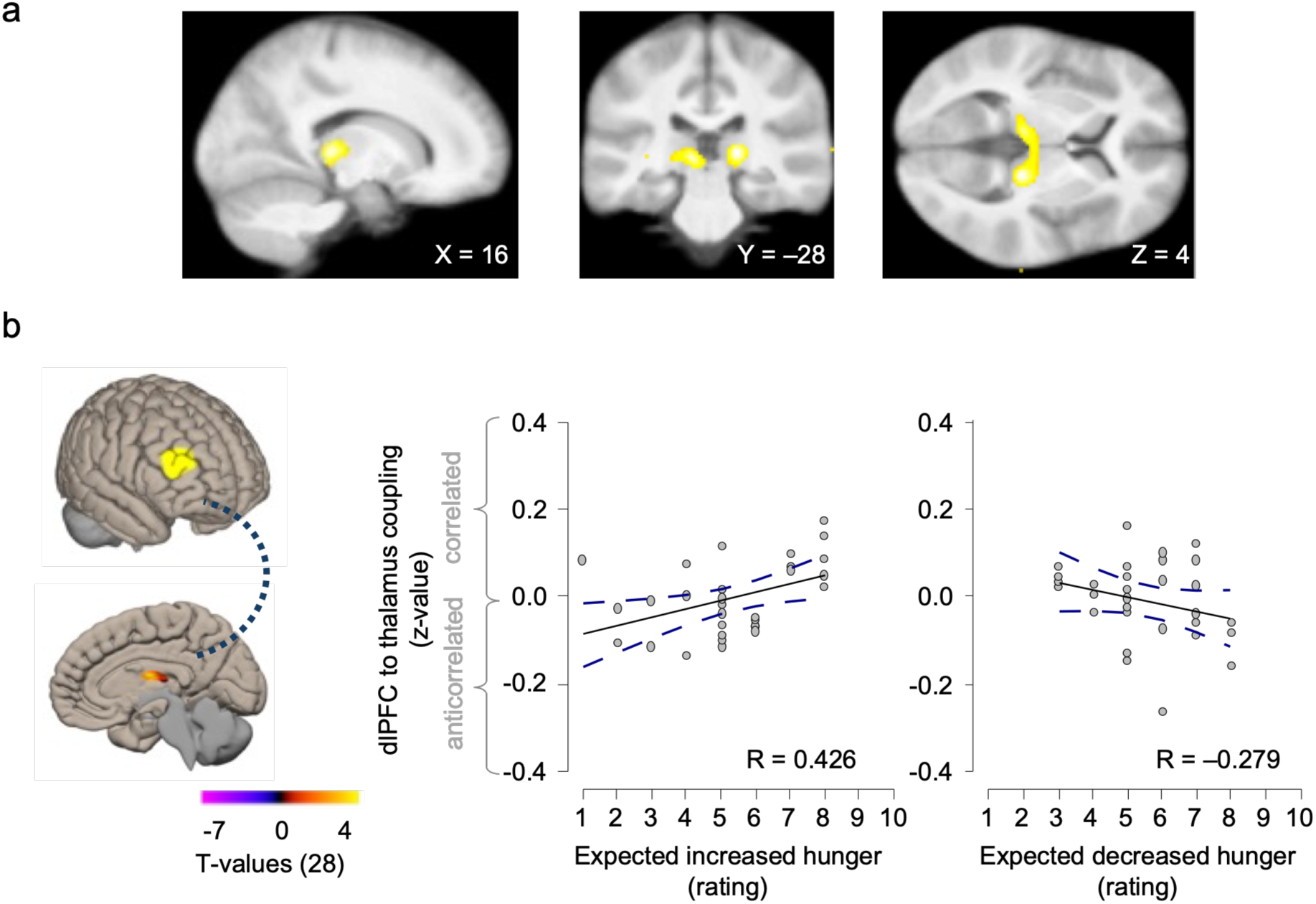
Difference between increased (n=30) and decreased (n=30) hunger suggestion groups in the dlPFC – to – thalamus connectivity moderated by expectation ratings (RS-fMRI full sample of n=60). (a) SPMs display the direct contrast of increased > decreased hunger expectations of the seed region, the dlPFC to the thalamus, at a family-wise error-corrected threshold of pFWE < 0.05 at the cluster level. They are displayed at a whole-brain height (voxel) threshold of p < 0.001 uncorrected and superimposed on the average anatomical brain. (b) Seed dlPFC and thalamus clusters are further displayed superimposed on a 3-D brain. The scatterplots show individual participants’ hunger expectations and z-values extracted from individual thalamic peaks selected via a leave-one-out procedure. R – Pearson’s correlation coefficient

## DISCUSSION

In this study, we used a placebo intervention targeting hunger sensations to induce hunger expectations and tested the effects of induced expectations on brain activity at rest. We leveraged a theory-driven approach and focused on the seed-to-voxel connectivity of three regions of interest in the prefrontal cortex: the medial (mPFC), ventromedial (vmPFC), and dorsolateral (dlPFC) prefrontal cortex.

Their overall whole-brain connectivity was consistent with evidence that brain regions that co-activate at rest form distinct resting-state networks (Buckner et al., 2008). The mPFC and vmPFC were primarily connected to regions of the default mode network (DMN), of which they are also components (Raichle et al., 2001). The dlPFC was mainly connected to regions within the frontoparietal network (FPN), of which it is a component (Duncan, 2010). Importantly, participants’ expectations about the effectiveness of a placebo drink in reducing or enhancing hunger moderated this connectivity. Decreased hunger expectations led to reduced functional connectivity between the mPFC and angular gyrus, while increased-hunger expectations were associated with reduced functional connectivity between the vmPFC and the dm/dlPFC. When directly comparing the two groups, we found that increased hunger expectations led to stronger functional coupling between the dlPFC and the thalamus, and this pattern was reversed for decreased hunger expectations.

Participants fasted overnight before the study and reported feeling hungry from the start to the end of the experiment. The increase in hunger was more noticeable in the group suggested to have increased hunger than in the group suggested to have decreased hunger. This finding aligns with previous research involving a larger sample that included the RS-fMRI data from this study (Khalid et al., 2024). The observed moderation of functional resting-state connectivity between the prefrontal cortex and the rest of the brain by hunger expectations may link to the placebo effect on hunger sensations. The mPFC and the angular gyrus, as part of the DMN, are linked with action–outcome monitoring and conscious self-awareness (Raichle et al., 2001; Schacter et al., 2007). More precisely, the angular gyrus activates when expected and actual outcomes differ during goal-directed tasks (Van Kemenade et al., 2017; Zwosta et al., 2015), and the mPFC was activated, in the same sample, with decreased hunger expectations during food choices (Khaldi et al., 2024). A decoupling of the angular gyrus from the mPFC at rest may thus reflect the behavioral observation that participants who expected the drink to decrease their hunger reported lower, expectancy-congruent hunger sensations. Additionally, the decreased resting-state connectivity between the vmPFC and the dm/dlPFC in participants who expected the drink to increase hunger might reflect a relaxation of efforts to control food cravings or a greater alignment between hunger expectations and sensations. Both brain regions activate under self-controlled food choices (Hare et al., 2011, 2009; Hutcherson et al., 2012) and related cognitive regulation of decision-making (Rudorf et al., 2014). Likewise, the finding that the dlPFC and the thalamus correlated positively with increased hunger expectations and negatively with decreased hunger expectations may reflect the integration of internal hunger signals with higher-level beliefs about hunger. The thalamus processes interoceptive information (Critchley and Harrison, 2013; Jänig, 1996) and relays signals to the cortex for interpretation (Iwai et al., 2015). Through this relay, it helps regulate cortical activity and influences attention, executive function, and decision-making (Azzalini et al., 2019; Dehghani and Wimmer, 2019; Nakajima and Halassa, 2017).

However, RS-fMRI does not allow direct inference about underlying behavior or thoughts. The observed functional connectivity moderations by hunger expectations may not translate linearly into hunger sensations. Exploratory analyses linking task-related dietary self-control, or changes in hunger, to the observed seed-to-voxel connectivities did not yield conclusive results. In contrast, our findings identified the neural associations that paralleled the development of hunger sensations from baseline to the end of the experiment based on hunger expectations. More research is needed to unravel the role of resting-state brain activity when placebo-induced expectations stand at odds with actual interoceptive or exteroceptive signals. This research should provide more direct evidence to confirm these interpretations of our findings and to avoid reverse inferences (Poldrack, 2006). For example, evidence could stem from examining alternative hypotheses regarding the directionality of functional connectivity between these prefrontal, parietal, and subcortical brain regions, perhaps by investigating the temporal dynamics of brain activity under mismatch between expected and experienced sensations. Additionally, applying machine-learning-based brain markers, such as the Neurobiological Craving Signature (Koban et al., 2023b) and other related self-control and mentalizing signatures (Açıl et al., 2026; Koban et al., 2023a), could improve the prediction of internal states from observed brain activity.

Our findings stem from analyses focused on three a priori defined seed regions of interest located in the prefrontal cortex rather than on exploratory whole-brain searches. The prefrontal cortex seeds for this study were selected based on (1) their independent activation during a dietary decision-making task in the same sample (Khalid et al., 2024), and (2) their well-reported distinct roles in placebo effects on pain (see review Wager and Atlas, 2015). The convergence between our resting-state findings and the functional roles these regions play during placebo effects across domains supports, though not definitively, the reliability of observed effects. Nonetheless, replication in a larger, independent sample and without prior task-based fMRI is needed to confirm the robustness and generalizability of our findings. This is important because our sample size (n = 30 per hunger-suggestion group) is modest for detecting individual-differences (brain–behavior correlation) effects, which are known to require larger samples for stable estimation (e.g., Marek et al., 2022).

The study included only female participants, an a priori design choice motivated by evidence that women may be more susceptible to contextual effects on sensations, such as those generated by placebo interventions (Colloca et al., 2016; Olson et al., 2021; Theysohn et al., 2014). This choice also aimed to reduce sex differences in food behavior (Davy et al., 2006; Frank et al., 2010; Marino et al., 2011; Rolls et al., 1991), given the relatively modest sample size. While this approach strengthens internal validity by limiting a potential source of variance, it also restricts the generalizability of our findings to men and more heterogeneous populations. We cannot rule out potential confounders due to variation in menstrual cycles, hormonal contraception, or sleep quality. Future studies with larger and more diverse samples, including male and non-binary participants, and assessing hormones and cycles are needed to determine whether the resting-state connectivity patterns observed here reflect a sex-specific mechanism of placebo-induced hunger expectations or a more general feature of appetitive placebo responding.

In conclusion, our results provide evidence that placebo cues tied to hunger expectations influenced the intrinsic functional connectivity of the prefrontal cortex to brain regions associated with interoceptive and cognitive control processes. These correlational findings contribute to efforts to define and explain how incoming sensory evidence may become associated with higher-order prognostic beliefs about bodily states and thereby translate into bodily sensing. Moreover, identifying the neurofunctional sources for individual differences in expectancy-driven placebo effects on hunger sensations contributes knowledge relevant to identifying the neurocognitive targets in the fight against obesogenic environments.

## METHODS

### Ethical considerations

The study received ethical approval from the local ethics committee (Comité de Protection des Personnes EST III no 19.12.05). The research followed the Declaration of Helsinki. All participants have provided written, informed consent.

### Participants

Sixty female participants (mean age = 34.57 ± 1.74 years) were included in the resting-state fMRI study. They were recruited via a public advertisement in the Paris area. Only self-reported female participants were recruited to control for gender effects. The participants were screened for normal-to-corrected vision, no history of substance abuse or neurological or psychiatric disorders, no medications, right-handedness, and no metallic devices that could compromise MRI safety.

Participants were randomly assigned to two suggestion groups – a decreased-hunger suggestion group or an increased-hunger suggestion group. Groups were matched on age, level of education, and body composition (Table 4). To keep hunger-related variance small, all participants were tested in the morning between 8:00 am and 12:00 pm after overnight fasting; the last meal was no later than 9 pm the day before.

**Table 4.** Sociodemographics and body mass index full sample (n=60).

| **Hunger suggestion** | **Age** in years<br>(mean ± sem) | **Education**<br>(mean ± sem) | **BMI** in kg/m<sup>2</sup><br>(mean ± sem) |
| --- | --- | --- | --- |
| Decreased (n=30) | 33.8 ± 2.2 | 4.9 ± 0.4 | 24.5 ± 2.2 |
| Increased (n=30) | 35.3 ± 2.7 | 4.3 ± 0.4 | 23.8 ± 0.7 |
| Cohen's d<br>(decreased ≠ increased group) | -0.113 | 0.364 | 0.083 |

BMI – body mass index, sem – standard error of the mean. Education was measured in years after high school.

All participants received 60€ for their participation in the experiment.

### Placebo intervention

The placebo intervention consisted of a scripted administration of a glass of mineral water (see Khalid et al. 2024 SI for details). In short, for the two hunger-suggestion groups, the water bottle carried a label specifically designed to convey information about the ingredients as either decreasing or increasing hunger. In addition, the experimenter explained the label content, and all participants read an information booklet describing the ingredients and their purported effects on hunger (for further details on the placebo intervention (see Khalid et al., 2024, SI Section 7). The experimenter ensured that participants understood the instructions before pouring the water into a 25 cl glass (8.45 oz). Participants were instructed to drink the entire glass within approximately 2–5 minutes. Three female experimenters delivered the intervention, and they changed throughout the data collection period. Due to the labels, script, and booklets, blinding of experimenters was not possible.

### Hunger expectation ratings

Immediately after consuming the water, participants provided expectation ratings regarding the placebo drink’s effectiveness in decreasing or increasing their hunger (Figure 8). Expectations were assessed on a 10-point scale (1 = minimum), with participants in the decreased-hunger suggestion group asked how much they expected the water to decrease their hunger, and those in the increased-hunger suggestion group asked how much they expected it to increase their hunger.

**Figure 8.**
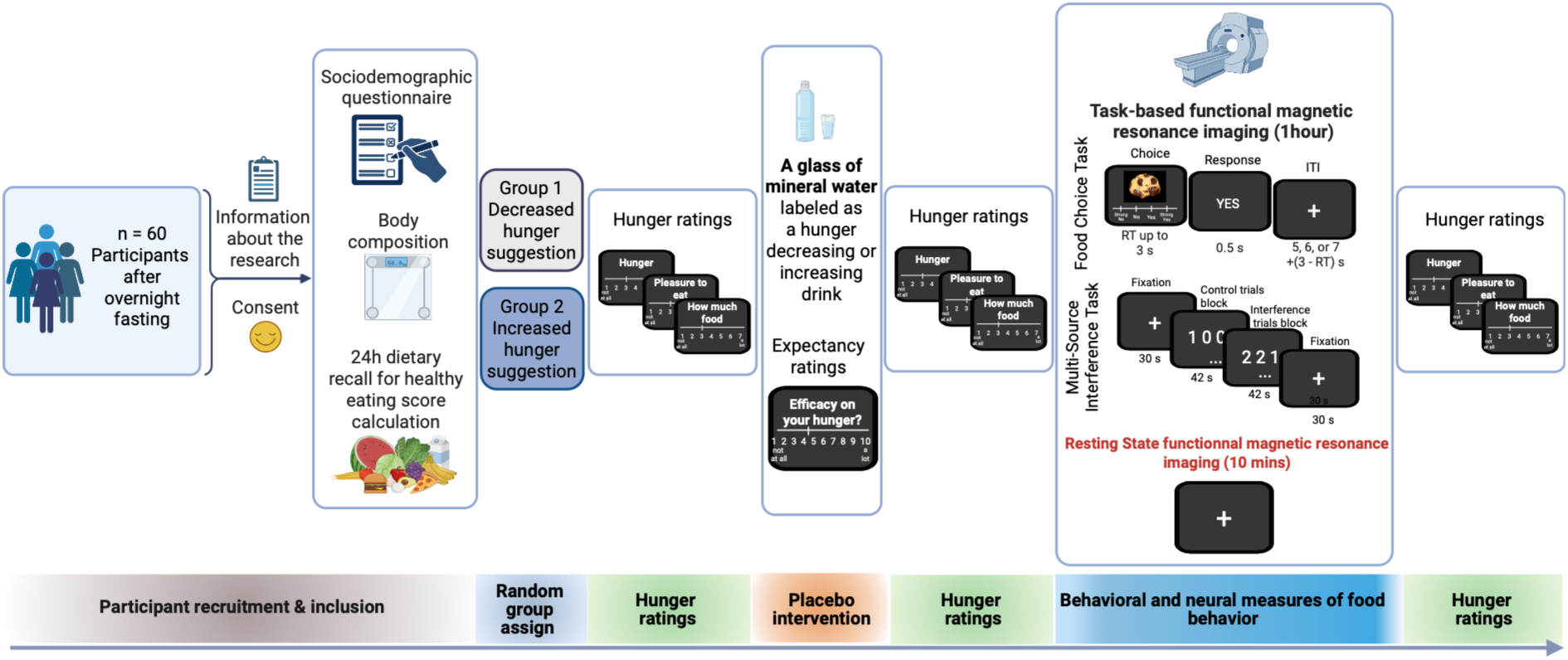
Overall experimental design for the RS-fMRU sample (n=60). Here, we focused on the resting-state fMRI component of the experiment. Task-based fMRI findings are reported in (Iraj Khalid et al., 2024).

### Hunger ratings

Subjective hunger ratings were obtained using three items: (1) overall hunger (‘How hungry do you feel?’), (2) homeostatic hunger (‘How much food could you eat right now?’), and (3) hedonic hunger (‘How pleasant would it be to eat right now?’). Each item was rated on a 7-point Likert scale (1 = not at all, 7 = very much), and responses were averaged into a composite hunger score. Ratings were collected at three time points: (1) baseline, (2) after the placebo intervention (post) but before the fMRI session, and (3) at the end of the experiment (Figure 8).

### MRI data acquisition

Resting-state activation was assessed after task-evoked brain activation for all participants. The fMRI session comprised four parts presented in a fixed order (Figure 8). First, a high-resolution structural scan was acquired for 5 minutes. This was followed by a 40-minute food-choice task (2x 20 minutes) and then by a 10-minute numerical version of the Multi-Source Interference Task (MSIT), which has been reported elsewhere(Khalid et al., 2024). The resting-state fMRI session was conducted at the end of the task-based fMRI sequences and lasted 10 minutes, during which participants were instructed to keep their eyes open and avoid falling asleep. They were told they could look at a fixation cross displayed on the computer screen.

T2-weighted multi-echo echo-planar images (mEPI) were acquired on a Siemens 3.0 Tesla VERIO MRI scanner equipped with a 32-channel phased-array coil. Three echoes were collected to optimize the trade-off between spatial resolution and signal quality in the orbitofrontal cortex (OFC)(Gowland and Bowtell, 2007; Poser and Norris, 2009). Each volume consisted of 48 axial slices acquired in an interleaved order. Acquisition parameters were as follows: echo times = 14.8, 33.4, and 52.1 ms; field of view = 192 mm; voxel size = 3 × 3 mm; slice thickness = 3 mm; flip angle = 68°; repetition time (TR) = 1.25 s. Whole-brain high-resolution T1-weighted structural images (1 × 1 × 1 mm) were obtained for all participants, co-registered with the mean mEPI images, and averaged to enable anatomical localization of functional activation at the group level.

### Preprocessing

Analyses of resting state fMRI data was performed using the CONN functional connectivity toolbox (RRID:SCR_009550 release 22.v2407) of the Statistical Parametric Mapping 12 software (SPM12, RRID:SCR_007037 release 12.7771) in Matlab (MATLAB_R2023b).

Prior to preprocessing, the three echo images per repetition time were summed into one EPI volume using the SPM12 Image Calculator and following a similar procedure reported in Khalid et al. 2024.

Preprocessing of resting-state time series involved realignment with correction for susceptibility distortion interactions, slice-timing correction, outlier detection, MNI-space normalization, and spatial smoothing with an 8 mm Gaussian kernel.

Potential outlier scans within each participant’s time series were identified using the ART toolbox in SPM using an intermediate setting of a framewise displacement above 0.9 mm and/or a global BOLD signal change above 5 standard deviations from a reference BOLD image. We computed the reference BOLD signal image for each participant by averaging all functional images, excluding motion-related images (see Table 5 for quality control overview).

**Table 5.** Quality Control (QC) of RS-fMRI time series.

| QC metric | Decreased Hunger suggestion (n = 30) | Increased hunger suggestion (n = 30) |
| --- | --- | --- |
| Mean FD (mm) | 0.10 | 0.12 |
| FD > 0.9 mm (%) | 1.96 | 1.75 |
| Mean number of valid volumes | 460 | 478 |
| Number of subjects passing QC | 25 | 26 |

FD – framewise displacement. Note: The 9 outlier subjects identified in the CONN toolbox's quality checks were included in the second-level analyses.

All participants, including those identified as outliers, were included in the statistical analyses to prevent unbalanced groups and avoid reducing the overall sample size. To account for movement artifacts and prevent inflated correlations, mean framewise displacement was incorporated into the second-level analyses as a covariate. Sensitivity analyses also showed that the RS-fMRI seed-to-voxel connectivity results remained consistent even when excluding the nine outlier participants.

### Denoising

Following preprocessing in SPM, the realigned, slice-timed, and normalized resting-state images were denoised using a standard denoising pipeline as implemented in the CONN toolbox(Whitfield-Gabrieli and Nieto-Castanon, 2012).

In more detail, we estimated and removed potential confounding factors from each voxel and subject using Ordinary Least Squares regressions. Confounding factors were non-neural signals of the white matter (5 regressors), and cerebrospinal fluid (5 regressors), head motion parameters and their first-order derivative from the realignment steps (12 regressors), and outlier scans identified by the ART toolbox (n < 173 images). Moreover, the regression removed variance explained by session effects and their first-order derivatives (2 regressors), as well as linear trends (2 regressors). The residual time series were then bandpass filtered between 0.01 Hz and 0.09 Hz.

Based on the number of noise terms included in this denoising pipeline, the effective degrees of freedom of the BOLD signal after denoising were estimated to range from 40.2 to 94.8 (average: 88.7) across all subjects. We did not regress out the global signal to avoid negative correlation stemming from global signal regression (Murphy et al., 2009). We then used the residual time series for statistical analyses.

### Behavioral analysis

Statistical analysis of behavioral variables was conducted using Jeffrey’s Amazing Statistics Program (JASP team, 2025; JASP Version 0.95.4). Four participants (2 in each suggestion group) were excluded from the behavioral analyses because they reported not being hungry at baseline (average hunger rating of < 2), which was an a priori exclusion criterion.

To assess whether the placebo intervention was effective, we compared the subsample of 56 hungry-at-baseline participants using two-tailed t-tests and Cohen’s d for effect sizes.

Hunger ratings were fitted by a linear mixed-effects regression that included the between-subject factor group and within-subject factor time (categorical factor: baseline/end-of-experiment) as fixed-effects regressors, following:

Hunger ratings = β0 Intercept + βGroup Group + βTime Time + βGroup x Time Group x Time+ (Intercept|Participant).

Post hoc paired and two-sample t-tests were used to further examine the main effects of group and time, as well as their interaction.

### Resting state imaging analyses

We focused on a theory-driven analysis following a hierarchical testing logic:

1. We tested how each seed-ROI connected to the rest of the brain, irrespective of hunger expectations.
2. We tested each seed-to-voxel connectivity for group-specific moderation by placebo-induced hunger expectations within each hunger suggestion group.
3. We tested the between-group differences by comparing how each seed-to-voxel connectivity moderation by hunger expectations varied between hunger suggestion group.

These analyses were implemented in the CONN Functional Connectivity Toolbox. To avoid reducing the sample size for RS-fMRI analyses, the 4 subjects who were not hungry at baseline and were excluded from the behavioral analyses were included in the RS-fMRI analyses. Excluding these 4 participants did not change the RS-fMRI results.

### Seed regions of interest

We employed a seed-to-voxel approach to examine the resting-state functional connectivity of three regions of interest. The medial prefrontal cortex (mPFC, MNI = [2, 58, 22]), which encoded hunger expectations during dietary decision-making(Khalid et al., 2024). The ventromedial prefrontal cortex (vmPFC, MNI = [0, 52, -12]), which encoded the value assigned to food stimuli during dietary decision-making(Khalid et al., 2024). The dorsolateral prefrontal cortex (dlPFC, MNI = [40, 42, 26]), which activated during interference resolution and connected to the vmPFC during dietary decision-making, indicating its role in self-control(Khalid et al., 2024). All three regions were defined as spheres of radius 5 mm centered at their coordinates in the Montreal Neurological Institute (MNI) space.

### Seed-to-voxel connectivity analysis

For each seed, voxel-wise connectivity maps were generated by correlating the residual BOLD signal of each seed region with those of all other brain voxels. This approach is particularly well-suited for exploratory hypothesis testing, as it adds a theory-based framework for where to search (Biswal et al., 1995; Fox and Raichle, 2007; Greicius et al., 2007). Functional connectivity was defined as the spatiotemporal covariance between the BOLD signal in voxels within the seed regions and the BOLD signal across all other voxels in the brain. Functional connectivity strength was represented by Fisher’s r-to-z transformation of the Pearson’s correlation coefficients assigned to each voxel in the brain. This led to three coefficient maps, one per seed region in each participant. The individual seed-to-voxel connectivity maps were then included in a second-level random-effects analysis with expectation ratings and mean framewise displacement as covariates. One-sample t-tests tested whether the connectivity maps were moderated by hunger expectations relative to zero within each hunger-suggestion group and for each seed region of interest. Two-sample tests with expectation ratings as covariates were used for the direct contrast of suggestion groups for each seed region of interest.

### Statistical thresholding of RS-fMRI activations

Significance of overall seed-to-voxel connectivity reported in the polar plots of Figures 2, 3, and 4 was inferred at a false discovery rate (FDR) corrected height (voxel) threshold of p_FDR_<0.05 and a family-wise error-corrected threshold of p_FWE_<0.05p_FWE_ < 0.05 on the peak and cluster levels (Chumbley et al., 2010). We also considered a more lenient uncorrected height (voxel) threshold of p < 0.001, with an extended threshold of k voxels to meet a cluster-level family-wise error-corrected threshold of p_FWE_ < 0.05, to examine how activations extended across the whole brain for visualization purposes and for activations reported in Tables 1, 2, and 3. Significant moderations of seed-to-voxel connectivity maps by expectations were inferred at a whole-brain cluster-level family-wise error-corrected threshold of p_FWE_ < 0.05.

Clusters were localized using the CONN-toolbox custom-built HCP Independent Component Analysis (ICA) network atlas, which contains 32 regions of interest grouped into 8 canonical resting state networks.

Individual z-values of functional seed-to-voxel connectivity were extracted using a leave-one-out procedure that included 60-second-level random effects analyses for each seed ROI, leaving out one participant at a time. The z-values were extracted for each left-out participant from 5-mm-radius spheres centered at the peak MNI coordinates showing the strongest moderation by hunger expectations in the training sample of 59 participants. The resulting 60 left-out z-values were then correlated with expectation ratings and displayed in the scatterplots shown in Figures 5, 6, and 7. Post hoc Fisher’s r-to-z transformations were used to compare whether these correlations differed in strength between suggestion groups.

## Notes

### Competing Interest Statement

The authors have declared no competing interest.

### Summary of Updates

Methods and results with more stringent control of movement artifacts; revised figures and tables. More careful discussion of the findings.

### Funding

- Paris Region fellowship program: 945298

